# Rigorous estimation of post-translational proteasomal splicing in the immunopeptidome

**DOI:** 10.1101/2021.05.26.445792

**Authors:** Kamil J. Cygan, Ehdieh Khaledian, Lili Blumenberg, Robert R. Salzler, Darshit Shah, William Olson, Lynn E. Macdonald, Andrew J. Murphy, Ankur Dhanik

## Abstract

Recently, *de novo* peptide sequencing has made it possible to gain new insights into the human immunopeptidome without relying on peptide databases, while identifying peptides of unknown origin. Many recent studies have attributed post-translational proteasomal splicing as the origin of those peptides. Here, we describe a peptide source assignment workflow to rigorously assign the source of *de novo* sequenced peptides and find that the estimated extent of post-translational splicing in the immunopeptidome is much lower than previously reported. Our approach demonstrates that many peptides that were thought to be post-translationally spliced are likely linear peptides, and many peptides that were thought to be trans-spliced could be cis-spliced. We believe our approach furthers the understanding of post-translationally spliced peptides and thus improves the characterization of immunopeptidome which plays a critical role in the immune response to antigens in cancer, autoimmune disease, and infections.

## INTRODUCTION

Antigen presenting cells use major histocompatibility (MHC) complexes I or II to present peptides to CD8+ or CD4+ T cells, respectively^1^. Characterization of the peptides presented to T cells, known as the immunopeptidome, is of major interest for the study of infectious disease, autoimmunity as well as cancer immunotherapy^2,3^. Identifying cancer associated peptides that illicit an immune response can point to safe and effective targets for cancer immunotherapy. The discovery and characterization of the immunopeptidome can be achieved using a multitude of technologies such as whole-exome sequencing, RNA sequencing, ribosome profiling and tandem mass spectrometry (MS/MS) based peptide sequencing. While next-generation sequencing approaches can characterize the potential immunopeptidome, only direct detection of peptides, like in MS/MS, can provide experimental evidence for the existence of peptides presented by MHC complexes. By combining information about potential and measured MHC-bound peptides, it is possible to reduce false-positives and accurately characterize the immunopeptidome^4–6^. When using next generation sequencing approaches, peptides sequences stemming from somatic missense and indel mutations can be determined. ^7,8,9,10^ Peptides can also originate from multiple other genetic and transcription-based aberrations, including cancer specific gene and transposon overexpression (e.g. cancer-testis genes, transposons, and human endogenous retroviruses (HERVs)), alternative splicing^11^, stop codon readthrough or upstream open-reading frame translation^12^.

Immunopeptidomics using peptide-MHC elution followed by MS/MS generally requires a reference database of potential peptides that might be detected. Recent advances in peptide spectra matching software allow omitting reference database searches and perform *de novo* sequencing, whereby the software identifies unknown peptides, post-translational modifications (PTMs) and amino acid substitutions directly from MS/MS. Using these methods, we can understand the diversity of peptides that may be bound to the MHC complex, but not their protein sources. For MHC I, the canonical mechanism of presenting peptides starts with proteasomal cleavage of proteins within the cytoplasm, generating fragments between 8 and 12 amino acids in length. Those peptides are then bound to the MHC I complex before its translocation to the cell membrane. However, some studies have suggested that, in addition to cleavage, proteasomes can catalyze the reverse reaction, ligating small peptides together in a process called proteasome-catalyzed peptide splicing (PCPS)^6,13–15^. While canonical cleavage generates peptides whose sequences are identical to the parental protein (herein called linear), pieces of spliced peptides can be from the same protein (herein called *cis*-spliced) or, theoretically, from different proteins (herein called *trans*-spliced) (**Supplementary Figure S1**)^6,15,16^.

Spliced peptides may be generated in a stochastic manner, making it challenging to generate MS/MS databases in which they could be found. Previous groups have used different approaches to solve this challenge. In Liepe et al., the authors developed a strategy to identify spliced peptides in the MHC-I immunopeptidome by mass spectrometry^17^. They generated a database containing all possible *cis*-spliced peptides, allowing for MS/MS spectra to be queried for *cis*-spliced peptides. Authors reported that about 30% of p-HLA are short-distance *cis*-spliced peptides. The same group also developed a pipeline for mapping the MHC class I spliced immunopeptidome of cancer cells^15^. They suggested a substantial (~25%) portion of peptides can be mapped with high confidence to *cis*-spliced sequences in HCT116 and HCC1143 cell lines derived from colon and breast carcinomas, respectively. *trans*-spliced peptides were excluded from analysis in Leipe et al., since their occurrence *in vivo* is controversial, and their addition to a database would massively increase its complexity. Later, Faridi et al. developed a bioinformatics workflow to identify linear, *cis*-, and *trans*-spliced peptides called hybridfinder. Hybridfinder first searches for exact matches of peptides in the UniProt human protein sequence database, then it searches for all possible *cis*- then *trans*-splice forms of that peptide in the human proteome (**Figure S2A**). The authors used hybridfinder to analyze MS/MS data consisting of peptides eluted from MHC I complexes purified from seventeen HLA-monoallelic cell lines. They reported that *cis*- and *trans*-spliced peptides represent up to 45% of MHC-bound peptides^6^.

The above studies argued that the large proportion of MHC-bound peptides that were not found in the human UniProt proteome could be explained by proteasomal splicing. Here, we use simulated random peptide sequences to show that the complexity of possibilities when searching for *cis*- or *trans*-spliced peptides leads to extremely high false positive rates. Additionally, we evaluate other potential sources of *cis*- and *trans*-peptides and propose a novel pipeline that searches for sources of peptides from multiple sources, with the order optimized to minimize the false discovery of peptides.

## RESULTS

### Expanding the search for sources of non-canonical human peptides

While previous studies have estimated that up to 45% of MHC-bound peptides that do not map perfectly to the UniProt human proteome are the result of post-translational peptide splicing^6,15–17^, we hypothesized that unmapped peptides may also come from other sources. We developed a multi-step approach where we expanded the queried databases to include other likely sources. We searched for peptides from non-canonically translated regions of the human genome, i.e. peptides from regions annotated as non-coding. For this source, we used OpenProt^18^ which includes all open reading frames (ORFs) at least 30 codons long, which we supplemented with the rest of the human genome translated into six frames. We also included translations of known transcribed elements, including long non-coding RNAs (lncRNAs)^19^, micro RNAs (miRNAs)^20^ and HERVs^21^. Below, we refer to this combination of sources as the expanded human proteome database. In addition, we also hypothesize that unknown SNPs, missense mutations, or recurrent errors in either transcription, translation, or MS amino acid identification could generate peptides with a single mismatch to a sequence encoded in the human genome. We searched for mismatched peptides using BLAT to align *de novo* peptides sequences to the expanded human proteome database (**Figure 1**). Finally, some peptides may originate from other organisms, especially bacterial or viral sources. For these sources we searched for *de novo* peptide sequences in the BLAST^22^ database (**see Methods**).

**Figure 1.**
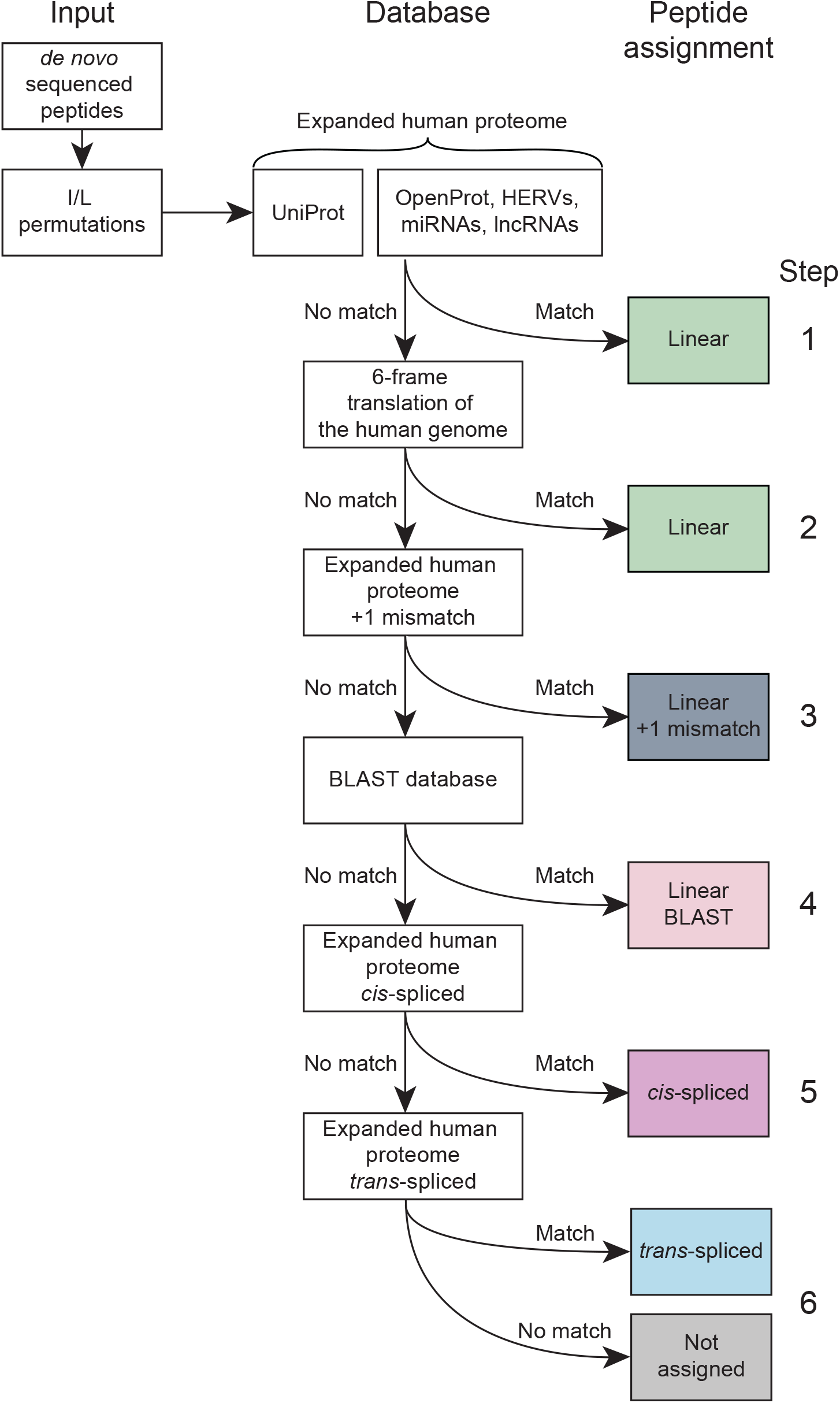
Detailed workflow of the peptide source assignment workflow.

### Estimation of False Discovery Rate

For each potential source of peptides described above (**Figure 1, databases**), including *cis*- and *trans*-splicing of peptides, we set out to estimate the chances of finding a match randomly. To estimate the false discovery rate (FDR) associated with each source, we asked how many randomly generated sequences could be found in each source. We generated 8-12mer peptide sequences (1,000 per length) in two ways: random sequences uniformly sampling all amino acids (referred to below as uniform random) (**Supplementary Table S1**) or sequences with frequencies of amino acids matching those found in vertebrates^23,24^ (referred to below as weighted random)(**Supplementary Table S2**).

We used the simulated sequences to estimate the FDR of each hypothesized sources of peptides (**Figure 2A**). Using the uniform random sequences, fourteen out of 5,000 peptides (0.28%) were found in the expanded human proteome (UniProt, OpenProt, lncRNAs, miRNAs and HERVs). When searching in canonically non-coding regions of the human genome using BLAT, 178 out of 5,000 (3.3%) of random peptides could be mapped. We also sought to estimate the FDR when searching for peptides mapping to the human proteome with a single mismatch; we found that 192 out of 5,000 (3.9%) peptides could be mapped. When searching for peptides that may come from non-human organisms in the BLAST database, 604 out of 5,000 (12.1%) sequences could be mapped. Finally, when we search for *cis*-spliced peptides in the expanded human proteome, 1,936/5,000 (38%) could be mapped; for *trans*-spliced peptides, 3,598/5,000 (71%) could be mapped (**Figure 2A**). For weighted random peptides, 50% and 68% of peptides could be assigned as *cis*- or *trans*-spliced, respectively (**Figure 2A**).

**Figure 2.**
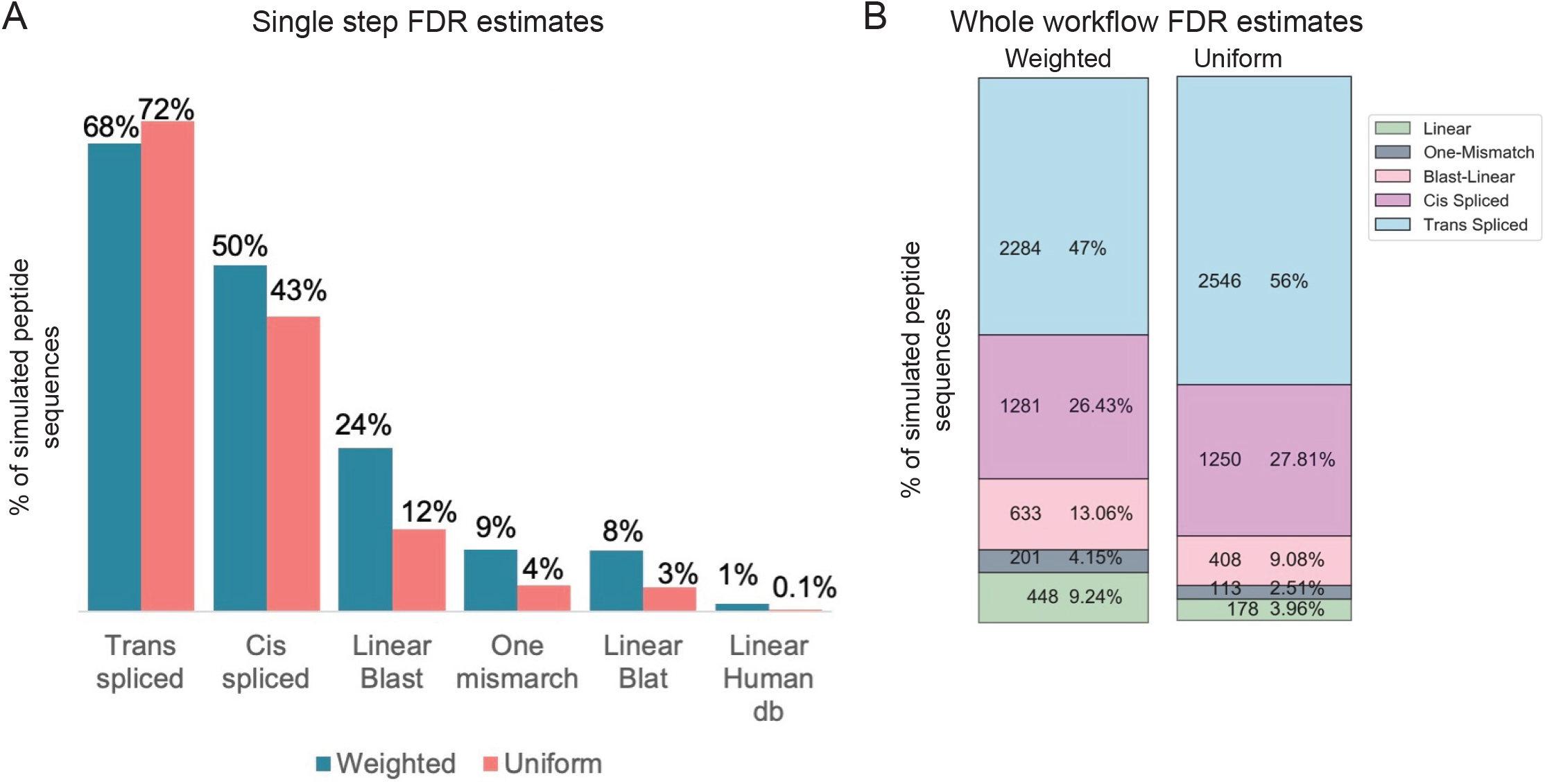
Estimating the false discovery rate of each putative source of peptides. a) Bar plot showing the percent of 5,000 randomly generated peptide sequences that could be matched at each step individually. b) FDR estimation of each step when the whole workflow is run in series.

We also estimated the FDR for the hybridfinder pipeline, which we recapitulated (**Figure S2A**). We found that none of the simulated peptides were assigned as linear, since hybridfinder only matches peptides to the UniProt database, not the expanded database we designed. However, for the uniform random sequence set, 772/5,000 (15%) could be assigned as *cis*-spliced and 2,329 of the remaining 4,228 (55%) could be assigned as *trans*-spliced. A total of 3,101/5,000 (62%) of uniform random sequences could be assigned as *cis*- or *trans*-spliced peptides (**Figure S2B**). For weighted random peptides, 78% of simulated peptide sequences could be assigned as *cis*- or *trans*-spliced (**Figure S2B**). This high FDR indicates that the search space of spliced peptides within the human genome is so large that post-translational splicing events in the immunopeptidome are likely over-estimated.

We set out to design an enhanced peptide mapping pipeline to assign sources for peptides in order of decreasing FDR. When using either set of simulated data to order peptide sources by FDR, the enhanced pipeline searches for peptide sources in the following order: 1) the expanded human proteome database (assigned as linear), 2) the non-coding regions of the human genome using BLAT (assigned as linear), 3) single mismatch peptides in the expanded human proteome (assigned as linear), 4) the BLAST database (assigned as linear), 5) *cis*-spliced and 6) *trans*-spliced peptides in the expanded human proteome. When ordered in series, 4,495/5,000 (90%) of uniform random sequences and 4,847/5,000 (97%) of weighted random sequences are found with this pipeline (**Figure 2B**); while this FDR is high, researchers can choose an appropriate threshold and exclude mapped peptides from high-FDR sources. Indeed, previous studies have excluded searches for *trans*-spliced peptides due to the presumed high FDR and assumed rarity of occurrence^15,25^.

### Re-analysis of monoallelic cell line data using the peptide source assignment workflow

We compared peptide identification by the peptide source assignment workflow versus hybridfinder on peptides eluted from MHC complexes on the data set from the hybridfinder publication: immunopeptidomics from a collection of cell lines engineered to express a single HLA allele^6^. For example, we found that for the cell line expressing HLA-A*02:04, of the 1,075 peptides classified as spliced by hybridfinder, 215 could be classified as linear from the expanded human database, 120 could be classified as linear with one mismatch, and 301 could be classified as linear from the BLAST database; overall 636/1,075 (60%) of putatively spliced peptides were reclassified as linear. Additionally, 133 of the peptides that were classified as *trans*-spliced could be reclassified as *cis*-spliced using an expanded human (**Figure 3A**). Across all cell lines, 36% of putative *cis*-spliced peptides could be reclassified as linear, and 45.9% of putative *trans*-spliced peptides could be reclassified as linear or *cis*-spliced (**Figure 3B**). The peptide source assignment workflow we present here shows that putative spliced peptides are likely peptides stemming from mutated DNA sequences, non-canonically spliced RNA sequences, non-canonically translated regions of the human genome, mismatched human sequences or bacterial proteins. Altogether, 20% of peptides are assigned as spliced peptides with the workflow presented here, down from 29% using hybridfinder (**Figure 3B**).

**Figure 3.**
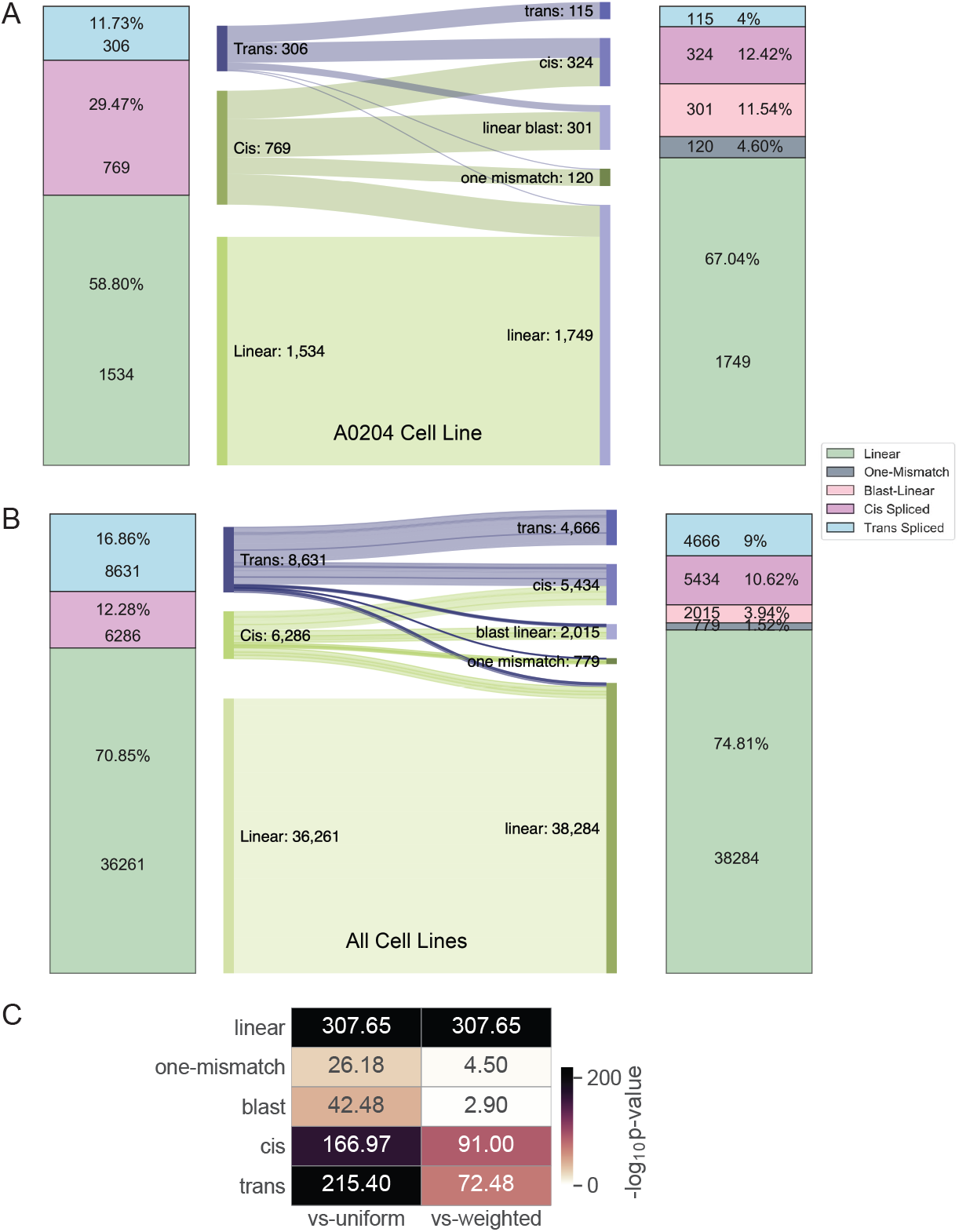
Peptide source assignment workflow results on peptides from HLA monoallelic cell line immunopeptidomics data. a) Peptide source identification on the HLA-A02:04-expressing the cell line by hybridfinder (left) and which peptides switched annotations (center) using the peptide source identification workflow presented here (right). b) Peptide source identification on all the HLA-monoallelic cell lines by hybridfinder (left) and which peptides switched annotations (center) using the peptide source identification workflow presented here (right). c) The Fisher’s exact test p-values measuring the enrichment of how many peptides were able to be assigned to a source at each step, compared to how many would be expected based on how many were assigned of the simulated random sequences.

At each step, we used the random peptides mapping results to estimate how many peptides were likely found by chance. When compared to weighted random peptides, more peptides detected in cell lines were assigned as linear (*P* < 1e-308, two-sided Fisher’s exact test), linear with a single mismatch (*P* = 3.16e-05), linear from the BLAST database (*P* = 0.00126), *cis*-spliced (*P* = 9.9e-92) and *trans*-spliced (*P* = 3.29e-73) (**Figure 3C**). More peptides were found that would be expected by searching for random peptides, this could indicate that each source is truly contributing to the immunopeptidome found in each cell line.

### Peptides that map throughout the human genome are enriched for expressed regions

We wanted to examine where peptides that map within the human genome, but outside of the UniProt proteome, land in terms of genomic annotations. The first three steps of the pipeline can map a peptide to regions of the human genome. In the first step, peptides that map exclusively in the OpenProt^18^ database land in ORFs that are not in the UniProt human proteome. We analyzed the location of these peptides in the human genome^26^ (**Figure 4A and Supplementary Table S3**), and found that, compared to the locations of all proteins in OpenProt, these peptides are enriched in exons, promoters and 5’UTRs (**Figure 4B**). The exonic enrichment is likely due to out-of-frame translation. In the subsequent step, peptides are mapped to a 6-frame translation of the human genome, though since OpenProt includes all proteins originating from ORFs longer than 30 amino acids, these peptides must come from ORFs shorter than 30 amino acids. The genomic annotation distribution of peptides mapped at this step is closer to that of the human genome, i.e. the majority of peptides map to intergenic or intronic regions (**Figure 4C and Supplementary Table S3**), possibly indicating these peptides assignments are contaminated with random matching. However, peptides from proteomic data sets were more enriched for exons, promoters and 5’UTRs than peptides from the random uniform or weighted simulations (**Figure 4D**). In the third step of the enhanced pipeline, peptides can be mapped to the expanded human proteome with a single mismatch (**Figure 4E and Supplementary Table S3**). Peptides mapped at this step show stronger enrichment in exons, introns and promoters than the enrichment found from peptides in the weighted or uniform simulated data sets (**Figure 4F**). At each step, there is consistent depletion of intergenic regions and enrichment of transcribed regions, as has been found in other studies focused on unidentified peptides in the immunopeptidome^27,28^. The enrichment of transcribed sequences supports the idea that the peptides assigned in these steps of the pipeline are correctly assigned, even though they do not map to proteins in the UniProt database.

**Figure 4.**
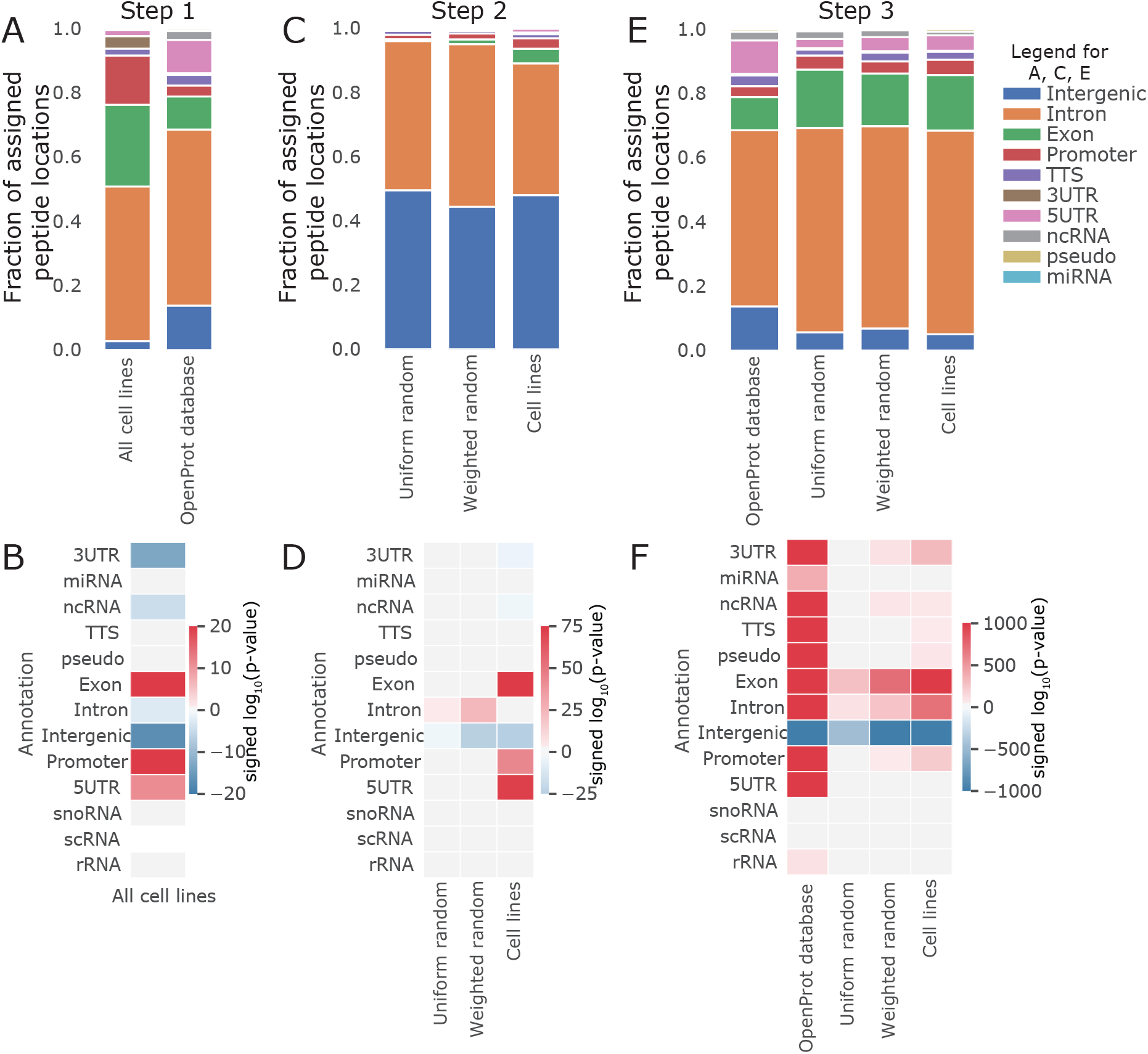
Genomic locations of peptides found outside of the canonical human proteome. a, c, e) Stacked bar plots showing the proportion of peptides that mapped to different genomic regions in steps 1 (a), 2 (c) and 3 (e) of the peptide source assignment workflow applied to the HLA monoallelic cell lines. b, d, f) Heatmaps showing enrichment of genomic annotations for the locations of mapped peptides in steps 1 (b), 2 (d), and 3 (f) of the workflow. b) signed log_10_ Fisher’s exact test p-values of the enrichment of peptides identified exclusively in the OpenProt database for the HLA monoallelic cell lines, versus the distribution of all proteins in the OpenProt database. d, f) signed log_10_ p-values calculated by HOMER, calculating enrichment of assigned peptide locations.

### Peptides identified by BLAST are not enriched for any microbial genus

The search within the BLAST database has the highest FDR for linear peptides in the peptide source assignment workflow. While peptides from cell lines had modestly more matches in the BLAST database than would be expected based on the uniform or weighted random data (**Figure 3C**), we determined if the BLAST assignments show enrichment of specific microorganisms which could be contaminants. To calculate enrichment, we removed peptides that could not be uniquely mapped to a single species; we then applied the Fisher’s exact test to the counts of peptides mapping to each genus in each cell line as well as all cell lines (**Supplementary Table S4 and Supplementary Table S5**). After correcting for multiple hypothesis testing, there were no significantly enriched genera in any cell line, or when considering all cell lines together. We hypothesize three possible, non-mutually exclusive causes for observing a lack of enrichment. First, there are either no contaminating organisms in the immunopeptidomics preparation. Second, the organisms present are not represented in the BLAST database. Third, the peptides stemming from contaminating organisms cannot be uniquely mapped to a single organism, and therefore were excluded from the above analysis. In the first two possibilities, it is not clear why the cell lines have more BLAST matches than expected. Taken together, these results do not support a biological underpinning for the peptides assigned at the BLAST step; rather, it is likely that the matches found at this step are random and spurious.

### Recurrent novel peptide

To identify reclassified common peptides across multiple datasets we selected peptides shared by more than three cell lines. For example, QSPVALRPL is highly recurrent, and was identified as *trans*-spliced by the hybridfinder algorithm, but was reclassified as linear using our pipeline. The same peptide is listed in the Immune Epitope Database as being a part of an unidentified protein. Upon further inspection, this is an out-of-frame peptide in the FAM96A gene, a pro-apoptotic tumor suppressor in gastrointestinal stromal tumors^29^. If out-of-frame translation is specific to cancer samples, this peptide could make an excellent cancer immunotherapy target.

## DISCUSSION

With the increasing number of peptides identified in immunopeptidomics experiments using *de novo* sequencing, the need for better characterization of the immunopeptidome is more pressing than ever. Previous studies attributed proteasome-catalyzed peptide splicing (PCPS) as the sole source of peptides of unknown protein identity. We present a peptide source assignment workflow that assigns parental proteins of *de novo* sequenced peptides from several sources with lower False Discovery Rate (FDR) than the set of all possible PCPS peptides. We found that 32% of putative PCPS peptides can be explained by single mismatches with known proteins, translational of supposedly untranslated parts of the human genome, or bacterial and viral peptides. Not surprisingly, the majority of peptides are encoded by known expressed regions. Finally, we identified a recurrent out-of-frame peptide in the tumor suppressor gene FAM96A that could be of interest as a cancer immunotherapy target.

Our pipeline has certain limitations; it is likely that a subset of peptides will be miscategorized in the pipeline described here. It is also possible that, by checking all possible permutations of leucine and isoleucine, due to their identical mass, we assign some peptides to incorrect proteins. We recommend that researchers incorporate expression and develop custom proteogenomic databases from RNA and whole exome sequencing to limit ambiguity of matches. In addition, as found in the data sets explored above, peptides found during the search of the BLAST step could all represent spurious matches, without evidence for a plausible source. In addition, there are additional sources of matches to *de novo* peptides sequences that, due to the complexity it would add to the search space, we have not added to our pipeline, such as PTM peptides. While there is undoubtedly some misidentification due to not accounting for PTM peptides, the addition of every possible PTM on every peptide would lead to too many possibilities to search.

Results from the peptide source assignment workflow presented here indicate that the extent of PCPS peptides may have been highly overestimated. The scope of the search space introduced when searching for cis- and trans-spliced peptides is enormous, and a large fraction of putatively spliced peptides can be explained by incomplete or inaccurate proteome database curation. While the contribution of PCPS is possibly overestimated in the past, our workflow annotates many more peptides as spliced than would be expected by chance. By checking other low-FDR sources first, we anticipate that the peptides that we annotate as PCPS are more likely to be enriched for actual post-translational spliced peptides. Importantly, While the total FDR of our workflow is extremely high, armed with the knowledge of the FDR of each step, researchers can filter out peptides assigned at steps with an FDR that is unacceptably high.

## METHODS

### Datasets

#### Simulated random peptides

We simulated two sets of peptide sequences for FDR estimation. We used the “random” built-in python library to produce sets of 8-12 length amino acid sequences, 1,000 peptides for each length, a total of 5,000 random peptides in each set. For the first simulated peptide sequence set, all amino acids have an equal probability of being incorporated into a sequence; we refer to this set as “uniform random”. In the second set, the amino acids have a probability of being incorporated that matches their frequency in vertebrates; we refer to this set as “weighted random”.

#### HLA-monoallelic immunopeptidomics

For the MS/MS data from HLA-I monoallelic cell lines, we downloaded the peptides from the supplementary table of Faridi et al. paper^6^. The data includes the expression of eight HLA-A alleles (A0101, A0203, A0204, A0207, A0301, A3101, A6802, A2402) and nine different HLA-B alleles (B5801, B5703, B5701, B4402, B5101, B0801, B1502, B2705, B0702). In total, there were more than 51,000 unique peptides.

### Recapitulation of hybridfinder

For ease of comparison to our peptide source assignment workflow, we recapitulated the workflow from hybridfinder^6^. First, each peptide is sought in the UniProt human reference proteome database. Peptides with perfect matches are annotated as linear. For peptides with no linear matches, we generate all possible splits of that peptide where the length of the smaller piece is longer than 1 amino acid. Then, we look for potential matches for each fragment through the database. We annotate the peptide as *cis*-spliced if we detect perfect matches of both fragments in a single protein. The matches can be reverse-ordered. Otherwise, if the matches are available in two distinct proteins, we annotate the peptide as *trans*-spliced. Peptides for which no split pairs match to any protein sequences are annotated as not available (N/A).

### The expanded human proteome database

We combined FASTA files of OpenProt, UniProt reviewed and unreviewed human sequences, which also includes protein sequences from some viruses that use humans as hosts (UniProt proteome version UP00000564^30^, downloaded in May 2020). We expanded this database to include translated proteins sequences from lncRNAs (NONCODE Version v5.0^19^, downloaded in May 2020), miRNAs (last modified 3/10/18, downloaded in May 2020), and endogenous viral elements (gEVE database ORFs^21^, downloaded in May 2020). We use this database when the workflow searches for linear human peptides and single-mismatched human peptides (steps 1 and 3), as well as in the search for cis- and trans-spliced peptides.

### The peptide source assignment workflow

We hypothesized possible sources from which peptides in immunopeptidomics experiments may be found, then measured the FDR inherent in each source using the simulated random datasets described above. We then ordered the steps of the workflow in order of ascending FDR to construct the workflow. The steps applied to each *de novo*-sequenced peptide are as follows:

*Step 1*: for perfect sequence matches in the expanded human proteome database (described above). Leucine (L) and isoleucine (I) have the same mass; therefore it is impossible to differentiate them in *de novo* search sequencing^31^. To account for this, for a given peptide containing I/L we considered all permutations of I and L residues. For example, for the peptide “ATTSLLHN” we have four possible permutations: ATTSLLHN, ATTSLIHN, ATTSILHN, and ATTSIIHN. If the algorithm finds a perfect match for any permutation, the peptides is annotated as “Linear”, and all possible protein sources of the peptide are included in the output. Otherwise the algorithm moves to step 2.
*Step 2*: Search for a perfect match in any of the six frames of the translated human genome using BLAT^32^. We used the following commands:

blat -t=dnax -q=prot -minScore=7 -stepSize=1 hg38.2bit Fasta_query output.psl
psl2bed < output.psl > perfect_match.bed If a perfect match is found, that peptide is annotated as “Linear” and possible source sequences are included in the output. Otherwise the peptide is passed to the step 3.
*Step 3*: Peptides are mapped to the expanded human proteome database, this time allowing one mismatch using this code:

blat -t=prot -q=prot -minScore=7 -stepSize=1 combined_DB.processed.fasta Fasta_query
output_blat_hits.psl” If a sequence with a single mismatch is found, the peptide is annotated as “one mismatch”. Otherwise the peptide is passed to Step 4.
*Step 4*: Sequences are mapped to other organisms using the BLAST NCBI tool. If any perfect matches are found the results are annotated as “Blast Linear”.
*Step 5*: For the remaining peptides, the algorithm generates all possible splits of the peptide where the length of the smaller piece is larger than 1. Then it looks for matches of both fragments in all human sequence databases. If there is a match for both chunks in the same protein, the tool annotates the peptide as “*cis*-spliced”. Otherwise, if there are hits for both fragments in two different proteins, the tool annotates the peptide as “*trans*-spliced”. The rest of peptides that do not have any matches are assigned as not available (N/A).

### Genomic location of BLAT hits analysis

Analysis of the genomic locations of BLAT hits was performed using the annotatepeaks.pl script from the HOMER suite^26^. Specifically:

annotatePeaks.pl ${file} hg38 -annStats ${file}.summary.txt

Only basic annotations were considered for further analysis. To calculate the enrichment of genomic locations of peptides found in the OpenProt database with either a perfect match (step 1) or with a single mismatch (step 3), we performed a fisher’s exact test to compare the number of peptides in each genomic annotation in the sample versus in the whole OpenProt database. For peptides that mapped to any translated region in the human genome (step 2), we used the p-value enrichment calculated by HOMER for over or underrepresentation of each genomic annotation.

### Tools

Python, bedops, psl2bed, BLAT, BLAST, HOMER.

## AUTHOR CONTRIBUTIONS

K.J.C., and A.D. conceptualized and designed experiments; K.J.C., E.K., L.B., R.R.S., and A.D. performed research; K.J.C., E.K., L.B., R.R.S., D.S., W.O., L.E.M., A.J.M., and A.D. analyzed data; K.J.C., E.K., L.B., and A.D. wrote the manuscript.

## SUPPLEMENTARY FIGURES

**Supplementary Figure 1.**
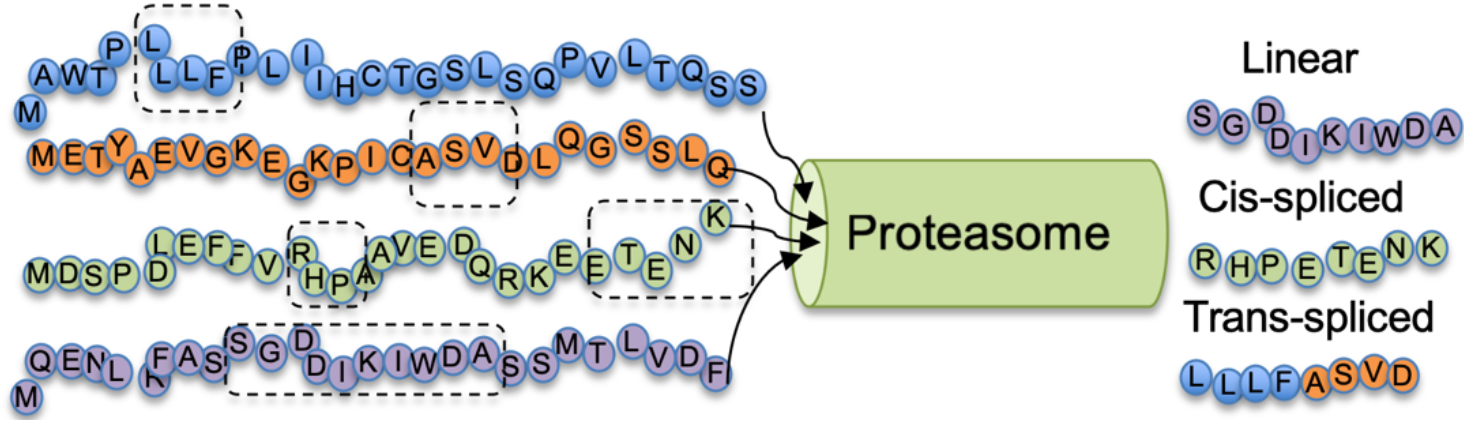
Linear, cis- and rans-spliced peptides made by proteasome-catalyzed peptide splicing/ligating. The linear peptide sequence matches perfectly to their parental protein, fragments of cis-spliced peptides are from the same protein, and the trans-spliced peptide fragments are from different proteins

**Supplementary Figure 2.**
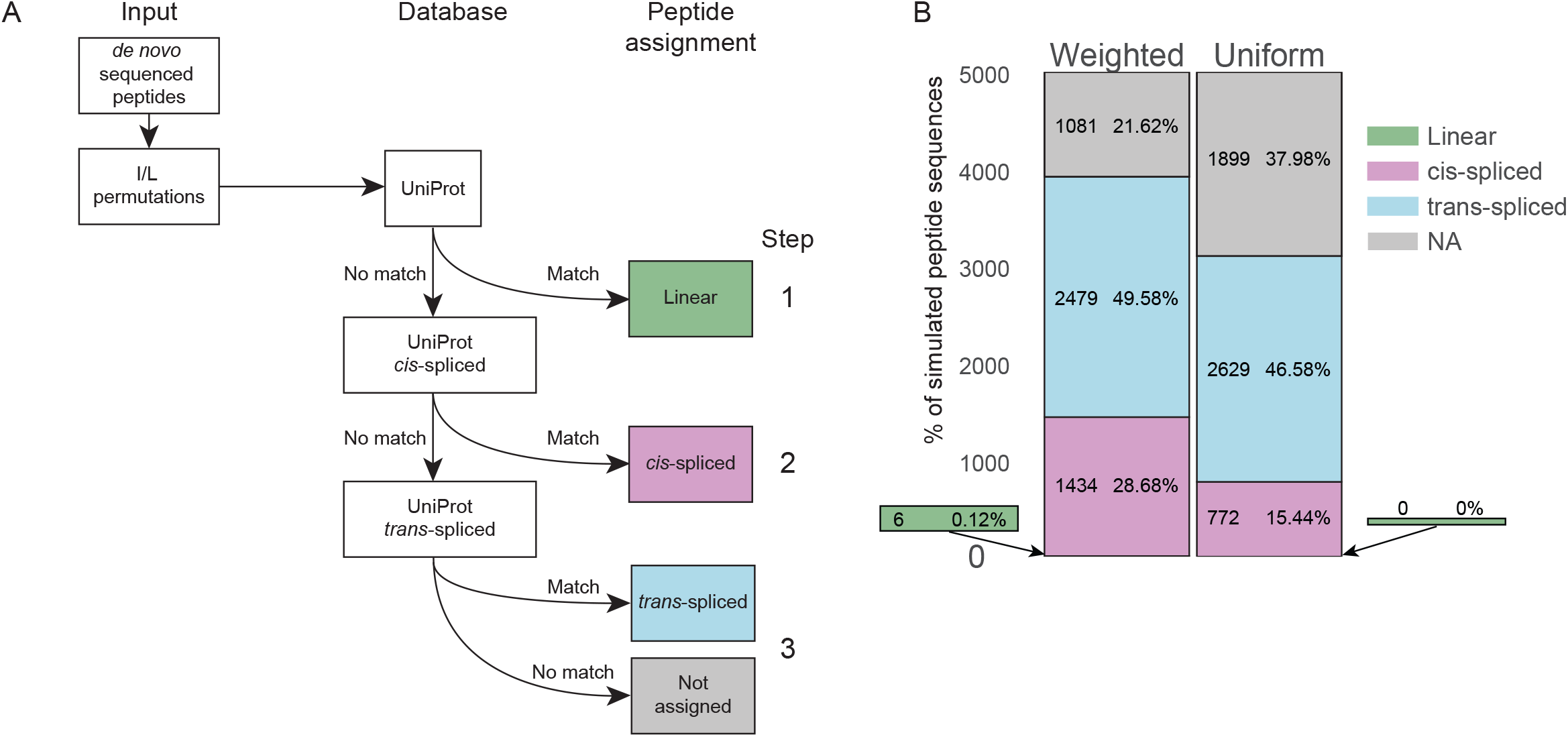
Recapitulation of hybridfinder. a) Schematic of the hybrid finder workflow. b) FDR estimation of the hybridfinder workflow using random sequences

